# Leakage-Aware Decoding of Music Perception and Cued Imagery Across the Full OpenMIIR EEG Cohort: A Reproducible Analysis of the Generalization Boundary

**DOI:** 10.64898/2026.08.06.743425

**Authors:** Yutian Wang, Keling Wang

**Affiliations:** School of Music, Georgia Institute of Technology, Atlanta, GA, United States of America; School of Music, Jiangsu Second Normal University, Nanjing, Jiangsu, China

## Abstract

Music perception and musical imagery provide a controlled setting for studying whether scalp electroencephalography (EEG) captures reproducible differences between externally driven and internally generated auditory states. We tested whether the public OpenMIIR dataset supports leakage-aware decoding of music perception versus cued musical imagery across its full ten-subject cohort, and we characterized the boundary beyond which the decoded signal fails to generalize. Using compact spectral and temporal EEG features, we applied stratified trial-grouped cross-validation, dummy and shuffled-label negative controls, leave-one-subject-out (LOSO) testing, a 1000-fold trial-level label-permutation test, and a group-level one-sided Wilcoxon test over per-subject within-subject accuracies, with Benjamini–Hochberg (BH) correction across the family of tested hypotheses. Within subjects, decoding was above chance at the population level: a group Wilcoxon test on logistic-regression accuracy gave *p* = 0.0195 with a large effect size (Cohen’s *d*_*z*_ = 0.95; 7 of 10 subjects above chance), confirmed by a pooled trial-level permutation test (*p* = 0.0040). Pooled trial-grouped balanced accuracy reached 0.567 [0.550,0.586] for random forest and 0.543 [0.523,0.561] for logistic regression, exceeding both dummy and shuffled-label controls. Cross-subject transfer was weaker and model-dependent: under LOSO, random forest reached 0.559 [0.523, 0.597], above its dummy baseline (uncorrected *p* = 0.014), whereas logistic regression did not generalize (0.518, *p* = 0.165). Under BH correction across the nine tested hypotheses, six comparisons survived at *q* < 0.05 (smallest *q* = 0.036), all involving the permutation test or the nonlinear model, while the linear model’s cross-subject contrasts did not. These results indicate that OpenMIIR EEG supports modest but reproducible within-subject discrimination of music perception and cued imagery, with a linear-within-subject versus nonlinear-cross-subject generalization boundary, and they show why public EEG music data require leakage-aware validation and calibrated subject-generalization claims.

## Introduction

Music perception and musical imagery engage auditory processing, memory, attention, affect, and reward-related mechanisms ^1^. They also offer a clean psychophysiological contrast: in music perception the auditory stimulus is externally present, whereas in cued musical imagery the participant internally generates or maintains a musical representation. Scalp EEG is well suited to studying broad temporal and spectral differences between such states, though it is poorly suited to claims about exact recovery of melody, harmony, timbre, or audio waveforms.

Public EEG music datasets make this contrast studiable without new human-subject data collection. OpenMIIR was designed for music imagery information retrieval and contains EEG recorded during music perception and imagination ^2,3^. Its value here is that the same musical material appears across heard and imagined conditions, allowing us to ask whether EEG features capture a broad perception-versus-imagery distinction under controlled validation. Closely related work by Ofner and Stober explored EEG representations of perceived and imagined music with a reconstruction-oriented ambition ^4^. Music-EEG research also frequently targets affective decoding, with datasets such as DEAP supporting emotion analysis from music-video stimuli ^5^, and reviews show that spectral EEG features and machine-learning classifiers can capture affect-related information under controlled conditions ^6^.

The present study is deliberately narrower than reconstruction or affective decoding. Prior OpenMIIR-related work established that the dataset is usable and motivated brain-to-music ambitions, but it did not systematically quantify, under leakage-aware validation and negative controls, *whether and how far* a perception-versus-imagery signal generalizes across the full cohort. That prerequisite question matters because EEG decoding results are easily inflated: when adjacent windows from the same trial leak across training and validation splits, when dummy baselines are omitted, when single-subject findings are generalized without subject-level testing, or when many model–control comparisons are reported without multiplicity control. Recent translational work emphasizes the practical impact of data leakage in EEG deep learning ^7^. For small public EEG datasets, validation design is therefore part of the scientific claim: a classifier that performs above chance only under window-level leakage, only within one participant, or only before multiple-comparison correction should not be read as evidence for a robust condition-level marker.

Compact neural architectures such as EEGNet are common EEG-classification baselines ^8^, and broader reviews show that convolutional and recurrent models can perform well but depend strongly on dataset size, preprocessing, and validation design ^9,10^. On small public datasets, conventional linear and tree-based models remain competitive because they impose fewer degrees of freedom and are less prone to overfit subject-specific structure. We therefore use logistic regression and random forest as primary, interpretable baselines.

This paper makes four contributions. First, it provides a leakage-aware reproducibility analysis of OpenMIIR perception-versus-imagery decoding over the *full ten-subject cohort* rather than a single participant. Second, it replaces small-fold screening statistics with confirmatory inference—a trial-level label-permutation test for the pooled model and a group-level Wilcoxon test over per-subject within-subject accuracies—with BH control across the comparison family. Third, it characterizes the generalization boundary by contrasting pooled within-subject decoding with leave-one-subject-out transfer, and shows that the boundary is model-dependent. Fourth, it distributes a reproducible workflow recording outputs, confidence intervals, negative controls, permutation nulls, and figures. Relative to prior OpenMIIR work ^2–4^, the novelty is not a new task or a reconstruction claim but a rigorous, cohort-level, validation-first characterization of what the perception-versus-imagery signal can and cannot support.

## Materials and Methods

### Dataset and Task

We use OpenMIIR EEG recordings and focus on binary condition decoding: music perception versus cued musical imagery. The analysis includes all ten OpenMIIR subjects with complete raw recordings (P01, P04, P05, P06, P07, P09, P11, P12, P13, P14). After event extraction and windowing, the dataset contains 8300 windows, 1200 physical trials, and 12 songs (Table 1). All raw EEG files were downloaded and verified for completeness against expected file sizes in an automated workflow before analysis.

Dataset summary for the full-cohort analysis.

Dataset summary for the full-cohort analysis.
| Property | Value |
| --- | --- |
| Subjects | P01, P04, P05, P06, P07, P09, P11, P12, P13, P14 ($n = 10$) |
| EEG channels (aligned) | 64 |
| Windows | 8300 |
| Physical trials | 1200 |
| Songs | 12 |
| Window length / hop | 3.0 s / 1.5 s |
| Conditions | music perception vs. cued musical imagery |

This is a retrospective analysis of an existing, publicly archived dataset. The OpenMIIR recordings were obtained from their public distribution and accessed for research purposes on 8 July 2026. The dataset is distributed in de-identified form, and the authors had no access to any information that could identify individual participants at any point during or after data collection; no attempt was made to re-identify participants. No new participants were recruited and no new recordings were collected for this study.

Raw EEG was read from the OpenMIIR FIF files with MNE. EEG channels were selected while excluding EOG, external (EXG), and stimulus channels. Because a subset of subjects (P09, P11, P12, P13, P14) contained additional external electrodes (EXG5/EXG6) absent for the others, all recordings were aligned to a common set of 64 EEG channels defined by the first subject; this alignment is what allows the full cohort to be pooled rather than silently dropping the affected subjects. Signals were resampled to 125 Hz when necessary and z-scored channel-wise before windowing. Trial events were read from the STI 014 channel and parsed with the OpenMIIR event-code convention, in which the trial code is stimulus ID multiplied by 10 plus condition ID. The binary task used condition 1 for music perception and condition 2 for cued musical imagery. EEG windows were 3.0 s long with a 1.5 s hop and were generated only within each trial’s duration.

### Feature Extraction and Models

EEG windows are represented with compact bandpower and time-domain features. For each channel, the extractor computes log-transformed mean spectral power in theta (4– 8 Hz), alpha (8–13 Hz), beta (13–30 Hz), and gamma (30–45 Hz) bands using the real-valued fast Fourier transform, and appends channel-wise time-domain mean and standard-deviation features. These features are interpretable and stable for small public EEG datasets. The primary models are dummy baselines (most-frequent and stratified), logistic regression (standardized features, balanced class weights, 2000 maximum iterations), and random forest (300 trees, maximum depth 8, minimum leaf size 3, balanced-subsample class weights). Interpretation emphasizes whether performance survives dummy and shuffled-label controls, permutation testing, and group-level inference, not whether any single model is nominally best.

### Validation and Statistical Analysis

The main protocol uses stratified trial-grouped cross-validation so that windows from the same trial cannot appear in both training and validation partitions. Trial groups combine subject, condition, stimulus, repeat, and trial identifiers. The default is five folds, reduced automatically when fewer groups per label are available. Multi-subject transfer additionally uses leave-one-subject-out (LOSO) validation, excluding all windows of a held-out subject from training.

We report three levels of evidence. (i) *Descriptive and control comparisons:* balanced accuracy, macro F1, bootstrap 95% confidence intervals (CIs; 10,000 fold-level resamples), paired differences against the best dummy baseline, and shuffled-label negative controls (training-partition labels permuted, validation labels intact). Fold-level paired sign-flip tests accompany these but, given few folds, are treated as screening only. (ii) *A confirmatory permutation test:* for the pooled model, a trial-level label-permutation test with 1000 permutations, in which condition labels are permuted at the physical-trial level (never at the window level), the full leakage-aware cross-validation is re-run per permutation, and the one-sided empirical *p*-value is (#{null ≥ observed} + 1)/ (*n*_perm_ + 1). (iii) *A group-level population test:* per-subject within-subject trial-grouped balanced accuracies are tested against chance (0.5) with a one-sided Wilcoxon signed-rank test, reporting Cohen’s *d*_*z*_. This is the primary, pre-specified population claim. One subject (P04) scored exactly at chance; following the standard convention this zero difference is dropped by the signed-rank procedure, so the test is performed on the remaining nine non-zero differences while all ten subjects are retained in the descriptive summaries.

Finally, the *p*-values of the substantive model–control comparisons are corrected with the Benjamini–Hochberg (BH) false-discovery-rate procedure. The correction family comprises the nine tested hypotheses: for each scope (pooled, LOSO) and each classifier (logistic regression, random forest), the contrast against the best dummy baseline and the contrast against the shuffled-label control, plus the pooled permutation test. Contrasts of a dummy baseline against the best dummy baseline are excluded from the family because they compare a baseline with itself and return *p* = 1.0 by construction rather than testing a hypothesis; including such degenerate entries would inflate the family size and attenuate every other *q*-value without adding inferential content. We report both uncorrected *p*-values and corrected *q*-values, and provide a sensitivity check under the wider family in Section 3.4. Analyses used scikit-learn and SciPy; code and derived tables are provided (Data Availability Statement).

### Use of Generative AI Tools

Generative AI assistants (OpenAI Codex and Anthropic Claude) were used during this study. Their use spanned three areas. First, implementation: the assistants helped write and refactor the Python analysis package, including the feature-extraction, trial-grouped cross-validation, permutation-test, group-Wilcoxon, and Benjamini–Hochberg routines, as well as the plotting and packaging scripts. Second, statistical analysis: the assistants were used to implement and independently re-verify the confirmatory statistics reported here, including recomputing the false-discovery-rate correction and cross-checking every reported value against the saved experiment outputs. Third, manuscript preparation: the assistants were used to draft and revise manuscript text, tables, and figure captions. All experiments were executed by the authors on the authors’ own computing resources, and all analysis outputs reported in this manuscript derive from those runs rather than from model-generated content. The authors verified the analysis code, independently checked the reported numbers against the archived result files, reviewed and edited all generated text, and take full responsibility for the content of this publication.

## Results

### Population-Level Within-Subject Decoding

Within-subject decoding was above chance at the group level. Across the ten subjects, within-subject trial-grouped balanced accuracy for logistic regression had a median of 0.552 and a mean of 0.561, with 7 of 10 subjects above chance; a one-sided Wilcoxon signed-rank test against 0.5 gave *p* = 0.0195 with a large effect size (Cohen’s *d*_*z*_ = 0.95).

The trial-level label-permutation test on the pooled logistic-regression model gave an observed balanced accuracy of 0.538 against a null mean of 0.500 (null SD 0.015), with empirical *p* = 0.0040 over 1000 permutations (Table 2). Per-subject within-subject accuracies are shown in Table 3 and Fig 1.

Confirmatory group-level statistics. The group Wilcoxon test is the pre-specified population-level claim; the permutation test is a trial-level confirmation on the pooled model.

| Test | Model | Effect size | $p$ |
| --- | --- | --- | --- |
| Group Wilcoxon (within-subject BA vs. 0.5), $W = 40, 7/10$ above | logistic regression | $d_z = 0.95$ | 0.0195 |
| Group Wilcoxon (within-subject BA vs. 0.5), $W = 39, 6/10$ above | random forest | $d_z = 0.45$ | 0.1377 |
| Trial-level permutation (pooled, 1000 perm; obs. 0.538 vs. null 0.500) | logistic regression | + 0.038 | 0.0040 ($q = 0.036$) |

Per-subject within-subject trial-grouped balanced accuracy (logistic regression), which enters the group Wilcoxon test. Chance is 0.5.

| Subject | P01 | P04 | P05 | P06 | P07 | P09 | P11 | P12 | P13 | P14 |
| --- | --- | --- | --- | --- | --- | --- | --- | --- | --- | --- |
| Balanced acc. | 0.627 | 0.500 | 0.531 | 0.493 | 0.493 | 0.664 | 0.623 | 0.604 | 0.506 | 0.573 |

Main balanced-accuracy results for the full ten-subject cohort. CI denotes the bootstrap 95% confidence interval.

| Validation | Model | Balanced accuracy | Macro F1 |
| --- | --- | --- | --- |
| Pooled trial-grouped CV | random forest | 0.5667 [0.5495, 0.5863] | 0.5663 |
| Pooled trial-grouped CV | logistic regression | 0.5425 [0.5230, 0.5608] | 0.5424 |
| Pooled trial-grouped CV | dummy (stratified) | 0.5016 [0.4976, 0.5073] | 0.5015 |
| LOSO | random forest | 0.5587 [0.5234, 0.5973] | 0.4965 |
| LOSO | logistic regression | 0.5182 [0.5010, 0.5400] | 0.4708 |
| LOSO | dummy (stratified) | 0.5028 [0.4992, 0.5069] | 0.5026 |

**Fig 1.**
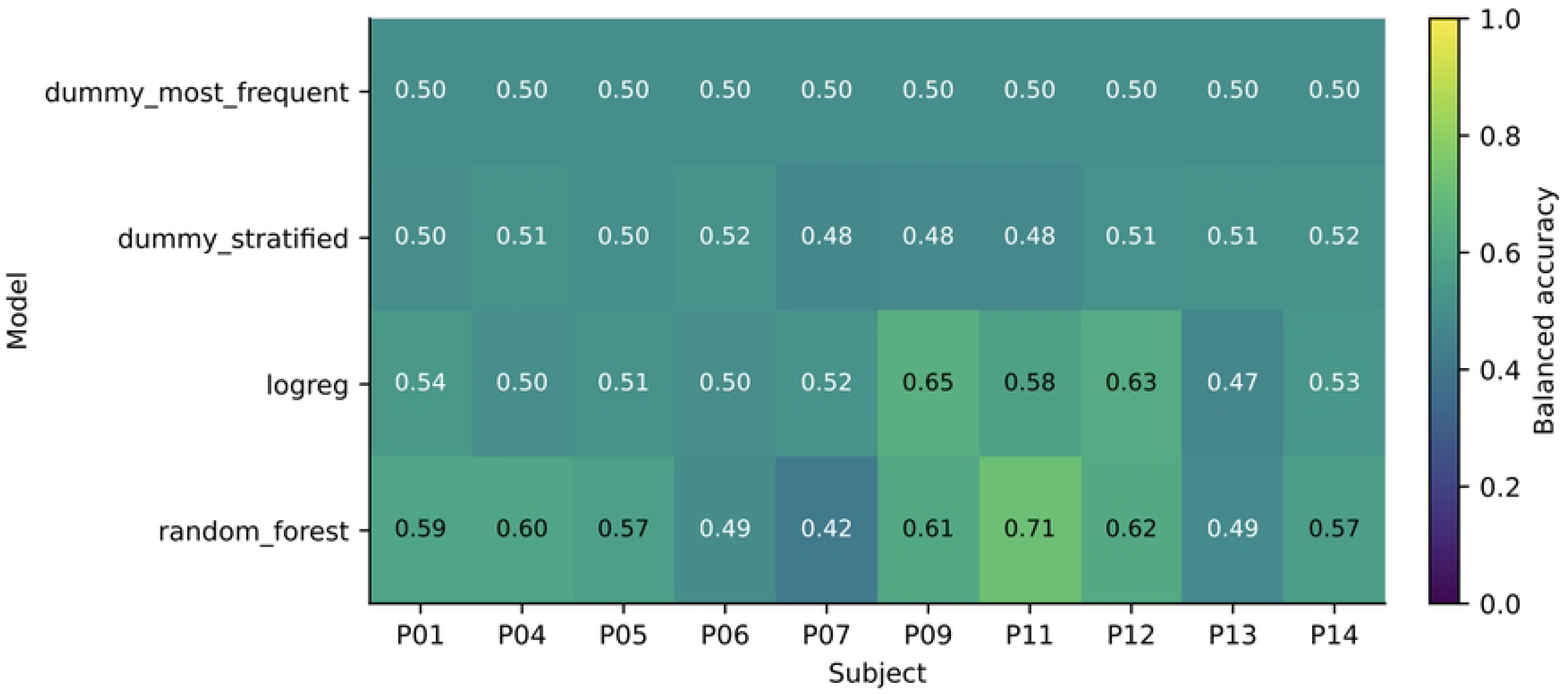
Per-subject decoding across the ten-subject cohort. Within-subject decoding is heterogeneous: several subjects are clearly above chance while a few sit at chance, consistent with the group-level test.

### Pooled and Cross-Subject Decoding

In pooled trial-grouped cross-validation, random forest reached a balanced accuracy of 0.5667 [0.5495,0.5863] and logistic regression reached 0.5425 [0.5230,0.5608]; both CIs exclude the dummy baselines, which sat at chance (0.5016 and 0.5000). Under LOSO, performance was lower and model-dependent: random forest reached 0.5587 [0.5234, 0.5973], a CI that excludes chance, whereas logistic regression fell to 0.5182 [0.5010, 0.5400] (Table 4, Fig 2).

Control comparisons (positive differences favor the normal-label model). p values are uncorrected fold-level sign-flip tests; q values are Benjamini–Hochberg-corrected across the family of nine tested hypotheses (these eight contrasts plus the pooled permutation test). Bold q values survive at q < 0.05.

| Validation | Model | Comparison | Difference | $p$ ($q$) |
| --- | --- | --- | --- | --- |
| Pooled CV | random forest | best dummy | $+0.0637$ | 0.0312<br>**(0.047)** |
| Pooled CV | logistic regression | best dummy | $+0.0395$ | 0.0625<br>(0.074) |

| Validation | Model | Comparison | Difference | $p(q)$ |
| --- | --- | --- | --- | --- |
| Pooled CV | random forest | shuffled labels | + 0.0622 | 0.0312<br>(**0.047**) |
| Pooled CV | logistic regression | shuffled labels | + 0.0420 | 0.0312<br>(**0.047**) |
| LOSO | random forest | best dummy | + 0.0551 | 0.0137<br>(**0.047**) |
| LOSO | logistic regression | best dummy | + 0.0146 | 0.1650<br>(0.165) |
| LOSO | random forest | shuffled labels | + 0.0429 | 0.0195<br>(**0.047**) |
| LOSO | logistic regression | shuffled labels | + 0.0233 | 0.0654<br>(0.074) |

**Fig 2.**
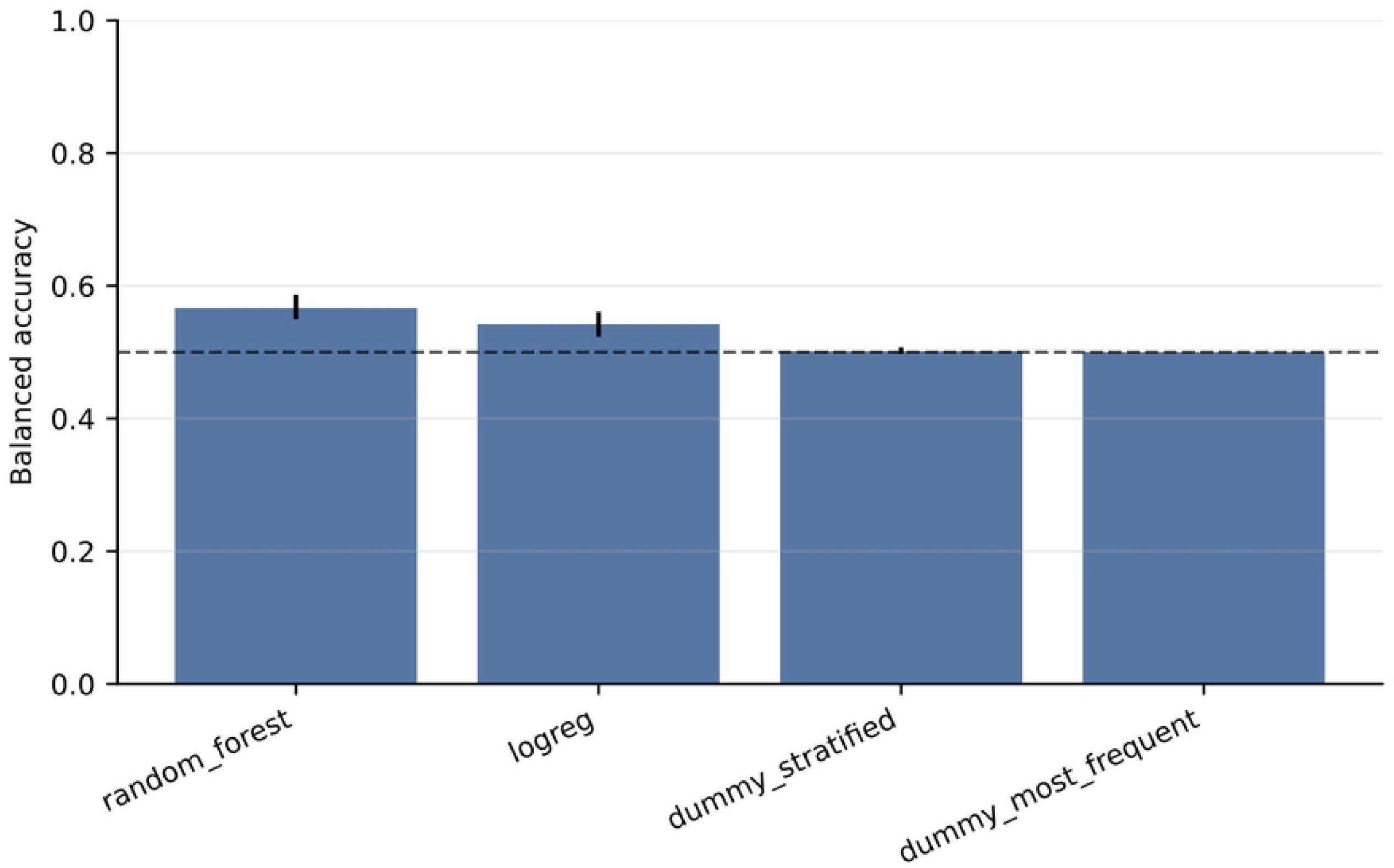
Pooled trial-grouped balanced accuracy for the full cohort. Error bars show bootstrap 95% CIs; the dashed line marks chance.

### Controls and the Generalization Boundary

In the pooled setting, both models exceeded the best dummy baseline and the shuffled-label control, with difference CIs excluding zero (Table 5). The generalization boundary appears under LOSO. Random forest transferred weakly but measurably to held-out subjects: its LOSO advantage over the best dummy was + 0.0551 [0.0183,0.0951] (uncorrected sign-flip *p* = 0.0137) and over the shuffled-label control + 0.0429 (*p* = 0.0195). Logistic regression did not transfer: its LOSO advantage over the best dummy was + 0.0146 [−0.0040,0.0373] (*p* = 0.165), with a CI including zero. The boundary is therefore *model-dependent*: linear features support within-subject decoding but do not transfer across subjects, whereas the nonlinear model captures a weak subject-independent component (Fig 3).

**Fig 3.**
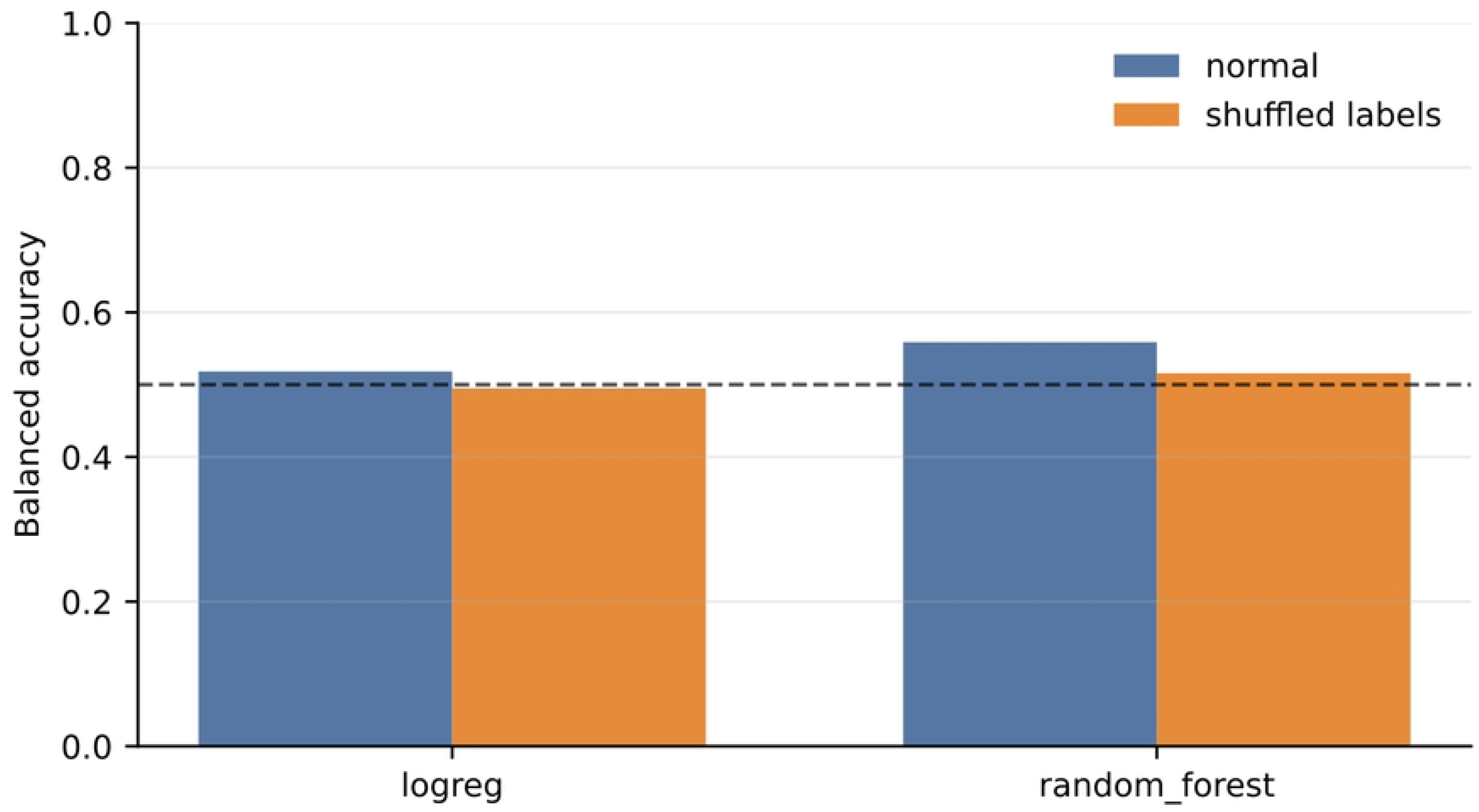
Leave-one-subject-out decoding versus the shuffled-label negative control. The nonlinear model keeps a positive normal-minus-shuffled margin across held-out subjects; the linear model does not clearly separate from its control.

### Effect of Multiple-Comparison Correction

Because several model × scope × control comparisons are reported, we applied BH correction across the family of nine tested hypotheses (Table 5, *q* column). Six comparisons survived correction at *q* < 0.05: the confirmatory pooled permutation test (*q* = 0.036), both random-forest contrasts under LOSO (*q* = 0.047), both random-forest contrasts in the pooled setting (*q* = 0.047), and the pooled logistic-regression contrast against shuffled labels (*q* = 0.047). The three comparisons that did not survive all involve the weaker linear-model contrasts: pooled logistic regression versus the best dummy (*q* = 0.074), LOSO logistic regression versus shuffled labels (*q* = 0.074), and LOSO logistic regression versus the best dummy (*q* = 0.165). This pattern mirrors the model-dependent generalization boundary described above rather than cutting across it.

As a sensitivity check, defining the family more conservatively by additionally including the four degenerate dummy-versus-best-dummy contrasts (each *p* = 1.0 by construction) inflates the family from nine to thirteen, scales every *q*-value by 13/9, and leaves the smallest at *q* = 0.052 with the random-forest contrasts clustered at *q* ≈ 0.068. The rank order, direction, and relative magnitude of every effect are unchanged; only the nominal threshold crossing differs. We report both so that the conclusion can be judged independently of how the family is drawn. In either case the population-level claim rests on the pre-specified group Wilcoxon test (*p* = 0.0195), which is a single test requiring no family correction, supported by the permutation test.

### Exploratory Interpretability

As an exploratory, single-subject analysis (P01), a feature-importance decomposition indicated contributions from beta, gamma, alpha, and theta features (feature-group importances 0.198, 0.187, 0.165, 0.157), with highest-ranked channels including TP7, P10, Iz, C4, Fp2, T8, O2, CP6, F1, FC3, P7, and Oz. These single-participant values are reported as feature-level evidence from a decoding model, not as cohort-level or source-localized effects, and are not part of the confirmatory analysis.

## Discussion

The results support a conservative but positive interpretation: across the full OpenMIIR cohort, EEG contains modest, reproducible information for distinguishing music perception from cued musical imagery within participants. Unlike a single-subject finding, this conclusion is supported at the population level by a group Wilcoxon test with a large effect size and by a trial-level permutation test on the pooled model. The interpretable feature analysis suggests that broad spectral and temporal EEG characteristics, rather than fine-grained musical content, drive the decoding.

The generalization boundary is the second contribution and is more nuanced than “performance drops with more subjects.” Within subjects, even a linear model decodes above chance; across subjects, only the nonlinear random forest retains a weak but measurable advantage over dummy and shuffled-label controls, while the linear model does not transfer. Psychophysiologically, the perception-versus-imagery contrast thus has both a subject-specific component that linear spectral features capture and a smaller subject-independent component that a more flexible model is needed to recover.

The multiplicity analysis both supports and bounds these claims. Under BH correction the confirmatory permutation test and every random-forest control contrast survive at *q* < 0.05, whereas the linear model’s cross-subject contrasts do not—the same asymmetry visible in the uncorrected results, which suggests the correction is separating signal from noise rather than uniformly suppressing it. The corrected values nonetheless sit close to the conventional threshold, and a more conservative family definition moves them just above it, so we anchor the population claim on the pre-specified group test rather than on any single corrected comparison. This validation-aware reporting is what small public EEG datasets require: above-chance decoding, above dummy and shuffled-label controls, under trial- and subject-grouped splits, robust to a permutation null, and stable in ordering under multiplicity control, but with effect sizes modest enough that the boundary conditions matter as much as the point estimates. Relative to earlier OpenMIIR work that established dataset usability and reconstruction ambitions ^2–4^, the present study contributes the missing prerequisite: a leakage-aware, cohort-level estimate of how far the perception-versus-imagery signal actually generalizes.

For future music-cognition and brain–computer-interface work, these findings suggest establishing robust cognitive-state decoding under subject-level validation and permutation testing before claiming richer music-representation decoding. Controlled, self-collected data with precise audio, MIDI, and event synchronization may be necessary for stronger claims about rhythm, phrase, timbre, and performance intention than the condition labels available in public data.

### Limitations

Several limitations apply. First, although the analysis now covers the full ten-subject OpenMIIR cohort, that cohort is small by machine-learning standards and the effect sizes are modest. Second, the features are compact and interpretable but not designed to recover fine-grained musical structure, and the interpretability analysis is exploratory and single-subject. Third, this is a secondary analysis of a public dataset, so stimulus timing, recording quality, and task design are constrained by the original OpenMIIR protocol. Fourth, although six comparisons survive false-discovery control, the corrected *q*-values sit close to the conventional threshold and shift above it under a more conservative definition of the comparison family; the population-level claim therefore rests on the pre-specified group test, and independent replication on additional datasets would strengthen it.

## Conclusions

This study provides a leakage-aware, full-cohort reproducibility analysis of OpenMIIR music perception versus cued-imagery decoding. Across ten subjects, within-subject decoding is above chance at the population level (group Wilcoxon *p* = 0.0195, *d*_*z*_ = 0.95; pooled permutation *p* = 0.0040, *q* = 0.036), with clean dummy and shuffled-label controls that survive false-discovery correction. Cross-subject transfer is weaker and model-dependent, present only for the nonlinear model. These results define a realistic, model-aware boundary for public-data EEG music-cognition decoding and motivate future controlled, synchronized EEG music experiments.

## Supporting information

### S1 File

#### Reproducibility supplement

Analysis code, notebooks, derived result tables, submission figures, permutation nulls, file manifests, and SHA-256 checksums that reproduce all summary analyses reported here. Openly archived at https://doi.org/10.5281/zenodo.21754264.

## Acknowledgments

During the preparation of this manuscript and the underlying analyses, the authors used OpenAI Codex and Anthropic Claude for analysis-code development, implementation and verification of the confirmatory statistics, and drafting and revision of the manuscript text. The authors have reviewed and edited the output and take full responsibility for the content of this publication. Details of how these tools were used are given in Section 2.4.

## Ethics statement

Ethical review and approval were waived for this study because it is a secondary analysis of the publicly available, de-identified OpenMIIR dataset and involved no new recruitment of human participants and no new data collection. The original OpenMIIR recordings were collected under the approvals reported by the dataset authors ^2^, who also obtained informed consent from the original participants.

## Data availability

The raw EEG data are from the public OpenMIIR dataset ^2^. The analysis code, notebooks, derived result tables, submission figures, permutation nulls, file manifests, and SHA-256 checksums required to reproduce all reported summary analyses are openly archived on Zenodo at https://doi.org/10.5281/zenodo.21754264. Raw EEG files and copyrighted audio stimuli are not redistributed; the raw recordings must be obtained from the original OpenMIIR distribution.

